# Sleep spindles enhance latent working memory representations

**DOI:** 10.64898/2026.06.26.734777

**Authors:** Sophia A. Wilhelm, Elkan G. Akyurek, Bernhard P. Staresina

**Affiliations:** Department of Experimental Psychology, Faculty of Behavioural and Social Sciences, University of Groningen; Neurobiology Expertise Group, Groningen Institute for Evolutionary Life Sciences, Faculty of Science and Engineering, University of Groningen; Department of Experimental Psychology, University of Oxford; Centre for Integrative Neuroimaging, Department of Psychiatry, University of Oxford

## Abstract

Working memory (WM) allows recently encountered information to be maintained over short periods, yet neural mechanisms supporting this short-term accessibility remain debated. Recent work suggests that WM representations can be maintained without persistent neural firing, in latent synaptic states often termed “activity-silent” memory. Such states can be reactivated by brief perturbations of cortical networks (“pinging”), providing a way to probe otherwise hidden representations. If such latent states rely on synaptic plasticity, they may be sensitive to sleep-dependent recalibration of cortical circuits. Here, we test whether sleep spindles, transient thalamo-cortical oscillations linked to synaptic plasticity, shape post-sleep accessibility of WM representations. Thirty participants performed a visual WM task before and after a daytime nap while high-density EEG was recorded. During the post-nap task, brief visual impulses were used to probe latent WM states, enabling multivariate decoding of memorised content. We show that longer spindle duration during prior NREM sleep predicts both improved post-sleep WM performance and higher-fidelity impulse-evoked decoding of item-specific representations, even when controlling for pre-sleep WM ability. These relationships were selective to spindles; slow-oscillation duration and other oscillatory metrics showed no corresponding effects. Together, our findings link sleep spindle dynamics to the neural readout of activity-silent WM and suggest that sleep-dependent synaptic recalibration optimises cortical circuits for subsequent, short-term information processing.

## Introduction

Working memory (WM) allows us to store and manipulate just-experienced sensory information for short periods of time (Baddeley, 2010). Despite its ubiquity, the exact maintenance mechanism of WM storage remains the subject of lively debate. The traditional view holds that select neurons maintain WM representations through sustained delay-period firing or increases in oscillatory EEG power (Curtis & D’Esposito, 2003; Funahashi et al., 1989; Sreenivasan et al., 2014). Yet, empirical work has presented inconsistent findings: both monkeys and humans can perform WM tasks even when delay-period spiking or oscillatory EEG power is minimal (e.g., Bae & Luck, 2018; LaRocque et al., 2012; Shafi et al., 2007; Watanabe & Funahashi, 2014). These inconsistencies inspired a synaptic model (often referred to as “activity-silent” WM), which posits that transient changes in synaptic weights could support the memory without continuous spiking (Mongillo et al., 2008; Stokes, 2015). This synaptic storage is attractive because it would reduce metabolic cost, shield WM contents from interference by placing them in states distinct from ongoing sensory coding (though see Wilhelm et al., 2026), and still permit rapid, context-dependent reactivation (Stokes, 2015). Computational work first fleshed out how such silent storage could work (Kozachkov et al., 2022; Mongillo et al., 2008; Mongillo & Tsodyks, 2024, 2026; Pals et al., 2020). More recently, converging empirical evidence has ensued: high-density single-unit recordings in macaque prefrontal cortex revealed coordinated ∼200ms “on-of” population switches during maintenance rather than continuous firing (Panichello et al., 2024). This points to a hybrid mechanism in which latent synaptic states maintain the memory, and brief bursts of spiking read it out when needed.

Despite the appeal of this account, a major challenge has been the fact that non-invasive methods in humans cannot directly assess these activity-silent states. To overcome this limitation, researchers have used an impulse perturbation approach (“pinging”), which provides a method to reactivate item-specific representations to levels detectable by EEG measurements. In turn, this allows researchers to infer activity-silent WM states (Figure 1A; e.g., Kandemir, Wilhelm, et al., 2024; Kandemir, Wolff, et al., 2024, 2024; Karabay et al., 2025; Wolff et al., 2015, 2017; Wolff, Jochim, et al., 2020; Wolff, Kandemir, et al., 2020). To achieve this “pinging”, a high-contrast task-irrelevant stimulus is presented during maintenance periods of WM to reactivate latent representations, thereby allowing those reactivations to be measured from the EEG via multivariate pattern analysis methods (Grootswagers et al., 2017; Haynes & Rees, 2006). Physiologically, short-term synaptic plasticity could be the mechanism underlying these silent states (Mongillo et al., 2008). Specifically, the assumption is that synapses engaged during encoding may remain transiently potentiated, possibly by residual calcium as a vestige of recent action potentials (Mongillo et al., 2008), and will therefore be more likely to engage in response to a high-contrast stimulus. Although “ping” responses may also reflect residual ongoing spiking or population states (Barbosa et al., 2021; Weng et al., 2026), the impulse-evoked window remains the most sensitive EEG readout of latent synaptic contributions to WM.

**Figure 1:**
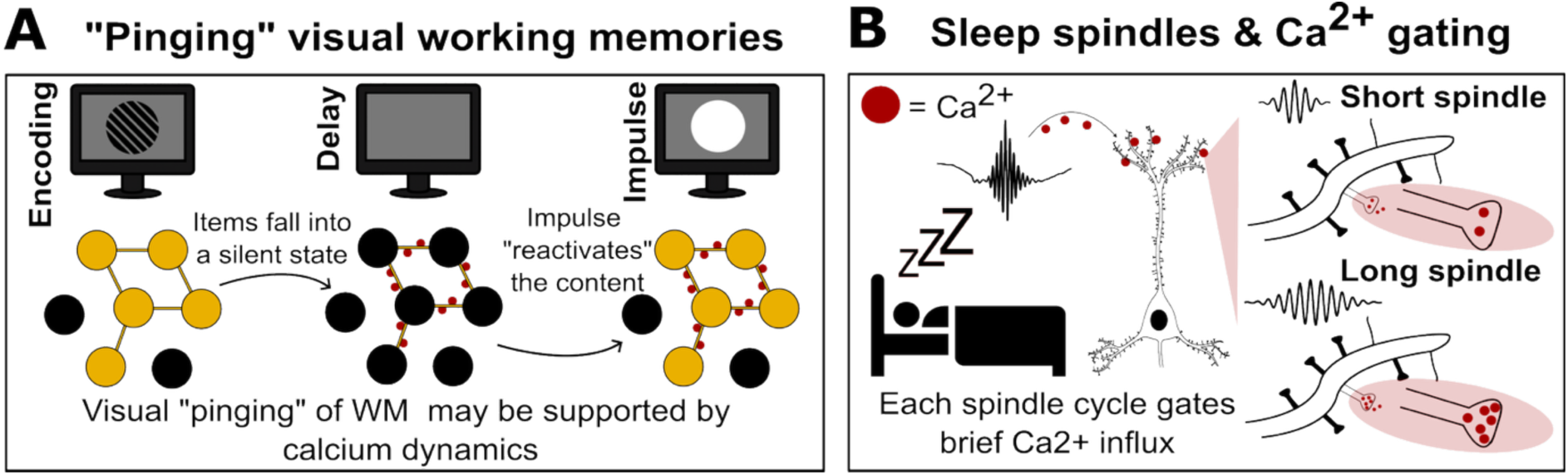
Hypothesised link between sleep spindles and subsequent “pinging” of activity-silent working memory. **A** In a classic visual working memory task, a grating is encoded through active neuronal firing, which is hypothesised to also lead to short-term synaptic changes (*left*). During the delay, this representation is maintained in an activity silent state (*middle*), presumably through synaptic plasticity. The residual calcium that keeps synapses potentiated is indicated by red dots. Finally, for EEG studies this information can be reactivated (and decoded) from its latent state using a brief, high-contrast impulse (“ping”) presented during the maintenance period (*right*). **B** During NREM sleep, spindle oscillations are accompanied by dendritic Ca2+ transients (depicted as red dots) in cortical pyramidal neurons. We propose that each spindle cycle gates brief Ca2+ influx at active synapses and that longer spindles, containing more cycles, provide an extended window for plasticity-related processes that tune synaptic weights. This spindle-driven synaptic tuning may determine how well latent WM representations can be decoded in a subsequent WM task.

Thus far, the impulse-perturbation (“pinging”) approach has largely been used as a stand-alone method to reveal WM representations that are not evident in ongoing delay-period activity. Here, we take a complementary approach. Rather than inferring activity-silent WM solely from the combination of weak or absent delay-period signals and successful impulse-evoked decoding, we ask whether these latent representations are sensitive to a physiological process known to reshape synaptic efficacy: sleep. Sleep is known to have profound effects on synaptic plasticity, supporting the strengthening of newly formed representations (Diekelmann & Born, 2010; Rasch & Born, 2013), while also engaging synaptic rescaling processes thought to restore network dynamic range and prepare the cortical circuits for subsequent sensory processing (de Vivo et al., 2017; Olcese et al., 2010; Tononi & Cirelli, 2006). If activity-silent WM relies on transient changes in synaptic efficacy, then sleep-related oscillations may alter the cortical substrate that determines how readily these representations can later be reactivated.

Among these oscillations, sleep spindles, i.e., waxing and waning thalamo-cortical rhythms in the 12–16 Hz range occurring during N2 and N3 sleep, have been identified as a key mechanistic vehicle for synaptic plasticity (Fernandez & Lüthi, 2020). At the cellular level, spindles are accompanied by dendritic Ca²⁺ influx in cortical pyramidal neurons (Seibt et al., 2017; Figure 1B), which can initiate molecular cascades involved in modifying synaptic efficacy and circuit function. Each spindle cycle coincides with brief Ca²⁺ bursts in thalamic neurons and propagates across the cortex, where most spindle events are spatially local rather than global (Andrillon et al., 2011; Piantoni et al., 2017). Although initiated in the thalamus, spindle duration is shaped by thalamo-cortical and cortico-thalamic interactions (Barthó et al., 2014; Bonjean et al., 2011). Together with evidence that the temporal profile of Ca²⁺ influx is critical for engaging plasticity-related pathways (Dittman & Regehr, 1998; Rosanova & Ulrich, 2005), this suggests that the duration of individual spindles may be especially relevant for driving synaptic recalibration: longer spindles, containing more oscillatory cycles, could provide extended windows for Ca²⁺-dependent tuning processes.

Building on this framework, we reasoned that spindle-mediated plasticity might influence the neural mechanisms underlying activity-silent WM. In synaptic models of WM, impulse-evoked decoding depends on the configuration of transient synaptic weights established during encoding. Sleep-dependent recalibration of cortical circuits could therefore shape how strongly and how selectively these latent states are reactivated after waking. When an impulse arrives, synapses that have been optimised during sleep (through selective strengthening, renormalisation, or improvements in local signal-to-noise) should generate a more robust and discriminable neural “echo” of the stored representation. We therefore hypothesised that individual differences in local spindle duration, as an index of cumulative Ca²⁺ gating during NREM sleep, would predict the fidelity with which latent representations can be decoded in a subsequent WM task. Consistent with this idea, behavioural work in individuals at clinical risk for psychosis has linked shorter spindle duration to poorer WM performance (Mayeli et al., 2022). Foreshadowing our results, we indeed found that longer spindle duration during a daytime nap predicted better WM performance, as well as higher fidelity in impulse-evoked decoding of activity-silent WM states, implicating spindle-driven synaptic recalibration as a candidate mechanism supporting post-sleep accessibility of latent WM representations.

## Results

The experimental protocol is outlined in Figure 2A. Participants (n = 30) came to the laboratory and were set up with high-density EEG. They first completed a visual WM task prior to sleep (Figure 2B), which served to index baseline, trait-like individual differences in WM performance. In this task, two memory items (circular gratings randomly chosen from 10 possible orientation angles) were presented sequentially, and a retro-cue indicated which of the two items participants had to respond to at the end of the trial. Responses were clockwise/counterclockwise judgements relative to the relevant memory item (rotation could differ across six angles). Average accuracy on this task was 81.1% (SD ± 7.7%), with an average response time of 682 ms (SD ± 86 ms).

**Figure 2:**
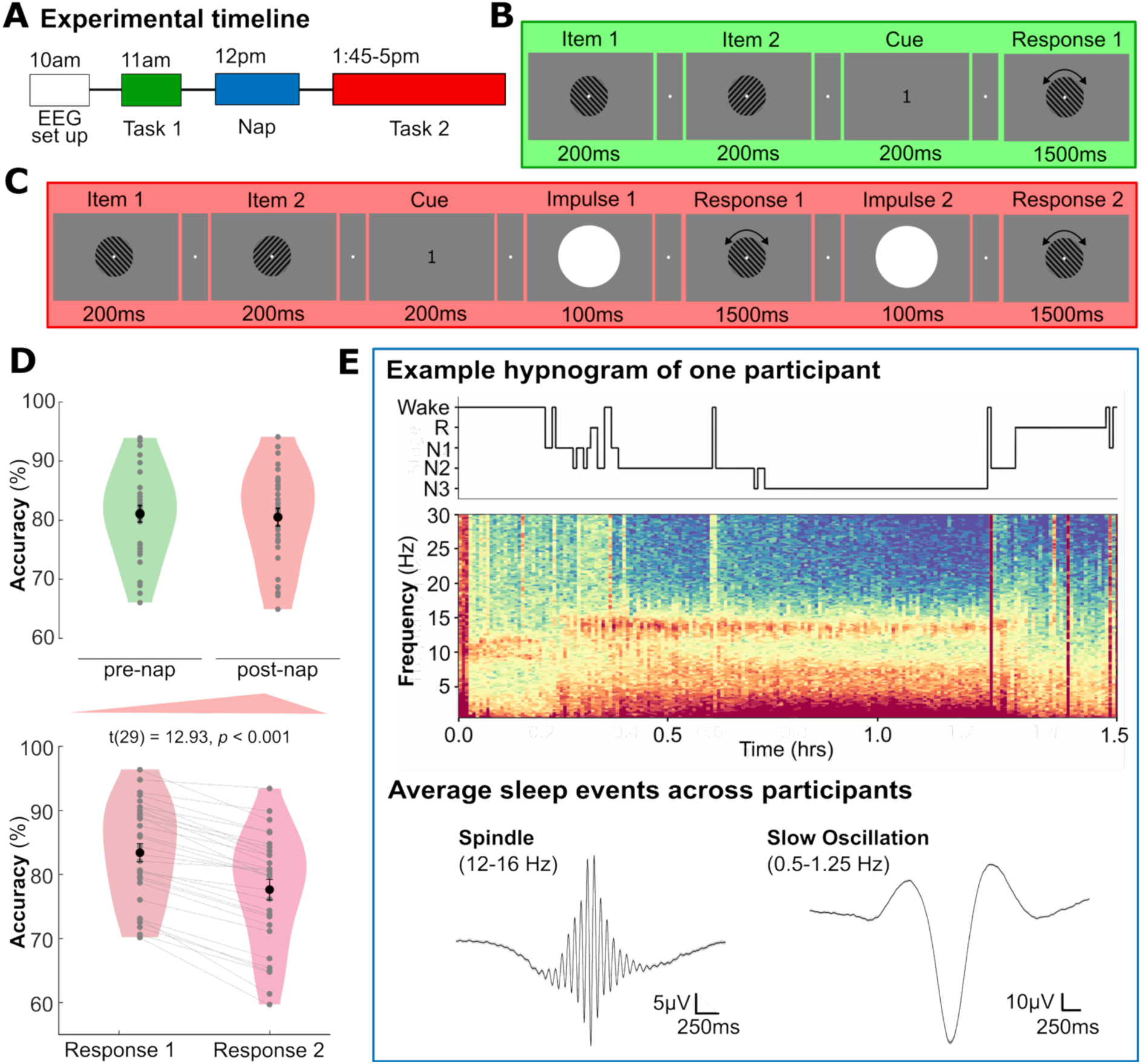
Experimental protocol, behavioural performance, and sleep oscillations. A Timeline of the experimental session. Participants (n = 30) were prepared for high-density EEG (58 channels), completed a pre-nap WM task (Task 1), underwent a 1.5h nap opportunity, and then completed a post-nap WM task (Task 2). B Task 1 (pre-nap) trial structure: Two oriented gratings (randomly drawn from 10 possible orientations) were presented sequentially (200ms each), followed by a retro-cue (200ms) indicating which item would be tested. Participants judged the probe as clockwise/counterclockwise relative to the cued memory item. C Task 2 (post-nap) trial structure: The encoding and cueing structure matched Task 1, but both items were probed on each trial in the cued order. To facilitate readout of activity-silent WM states, two brief visual impulses (“pings”, 100ms each) were presented prior to each response. D Behavioural performance. *Top*: Accuracy in Task 1(pre-nap) and Task 2 (post-nap). *Bottom*: Accuracy for Response 1 and Response 2 in Task 2 separately, showing a robust decline in accuracy for the second response (two-tailed paired t-test, *t*(29) = 12.93, *p* <0.001). Violin plots depict the distribution across participants; points indicate individual participants. Black markers denote group means + SEM. Grey lines show within-participant change. E Sleep measures during the nap. *Top*: Example hypnogram and corresponding time-frequency representation from one participant. Sleep spindles and SOs were detected during N2/N3 sleep. *Bottom:* Grand-average spindle (12-16Hz) at Cz, and SO (0.5-1.25Hz) at Fz.

Following this task, participants were given a 1.5 h nap opportunity. Average sleep duration during the nap was 67 min (SD ± 14.8 min), of which 47.9 min (SD ± 11.8 min) were spent in N2/N3 sleep (see Supplementary Table 1 for detailed sleep stages). After the nap, participants performed a second visual WM task (Figure 2C). The task was nearly identical to the pre-nap task; however, the retro-cue now indicated the order in which participants had to respond to the two memory items, as both items were probed on each trial. To facilitate the readout of activity-silent WM states via EEG, we included two brief visual impulses (“pings”), one preceding each response (Wolff et al., 2015, 2017). Mean performance in this task was 80.5% (SD ± 8.1%), with an average response time of 700 ms (SD ± 87 ms). For response 1, mean accuracy and RT were 83.4% (SD ± 7.8%) and 680 ms (SD ± 88 ms), respectively. For response 2, the corresponding values were 77.6% (SD ± 8.7%) and 710 ms (SD ± 90 ms), with a significant decrease in accuracy from response 1 to response 2 (t(29) = 12.93, p < 0.001, *d* = 2.36; Figure 2D). This decline is consistent with prior WM findings and is typically attributed to increased retention intervals and interference from retrieval of the first item (Barrouillet et al., 2004; Wolff et al., 2017).

During N2 and N3 sleep, we detected both sleep spindles and slow oscillations (SOs) in the nap EEG recordings (see Figure 2E for an example hypnogram; Supplementary Figures 2–3 for distributions and topographies of oscillatory metrics). We focused on spindle duration as our primary spindle metric of interest. To assess the physiological specificity of any observed relationships, we additionally quantified SO duration as a control rhythm.

### Sleep spindle duration predicts post-nap WM performance

As a first step, we asked whether mean spindle duration during the nap related to WM performance after sleep, and whether any such relationship could be attributed to baseline, trait-like differences in WM ability (i.e., pre-nap performance). To this end, we correlated spindle duration at each scalp electrode with mean task accuracy for each participant. No reliable association was observed between spindle duration and pre-nap (Task 1) performance (all p > 0.05, cluster-corrected; Supplementary Figure 4A). In contrast, spindle duration correlated positively with post-nap (Task 2) performance, yielding a significant centro-posterior cluster (cluster-corrected p = 0.015; Supplementary Figure 4C). This effect was selective to spindles: slow oscillation (SO) duration did not correlate with WM performance before or after sleep (both p > 0.05; Supplementary Figure 4B,D), and no other spindle or SO metrics showed significant associations (Supplementary Figure 4A–D).

To further corroborate the association between spindle duration and post-nap WM accuracy, we controlled for pre-nap performance using partial correlations. This analysis revealed a cluster over occipito-parietal sites in which spindle duration remained significantly related to post-nap (Task 2) accuracy (cluster-corrected p = 0.048; Figure 3A), consistent with a contribution of spindle physiology to post-sleep WM performance beyond baseline inter-individual WM performance differences.

**Figure 3:**
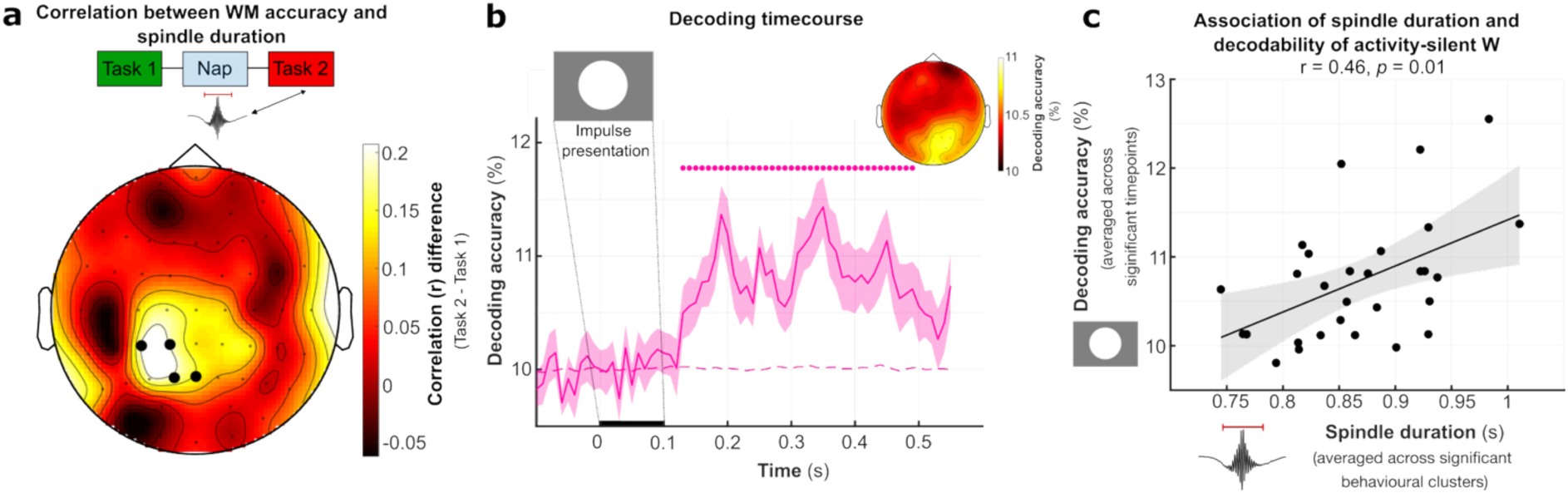
Sleep spindle duration predicts WM accuracy and impulse-evoked decoding of activity-silent WM content. **A** Partial correlation topography testing whether spindle duration predicts subsequent WM performance, controlling for pre-sleep WM accuracy. White dots indicate electrodes where higher spindle duration predicted subsequent WM performance (cluster-corrected *p* = 0.048). **B** Time course of orientation decoding accuracy (10-class LDA; 8-fold cross-validation) time-locked to impulse onset during the post-nap WM task. Solid line shows group-averaged decoding accuracy; the shaded area denotes ±SEM. Horizontal bar (pink) indicate the time window with significant above-chance decoding tested against a permutation distribution (cluster corrected *p* < 0.001). Dashed pink lines indicates 10% chance level. Inset topography shows the distribution of decoding accuracy across the scalp. Decoding accurac was obtained at each time point using a searchlight approach (including neighbouring electrodes as features). Decoding values were then averaged across the main time window of significance to obtain a single decoding accuracy value per electrode. **C** Across-participant association between mean spindle duration (averaged across the electrode cluster from Figure 3A) and decoding accuracy (averaged across significant time points from Figure 3B). Each dot represents one participant; the solid line shows the fitted regression line and shaded band indicates the 95% confidence interval (*Spearman’s rho* = 0.46, *p* = 0.01).

### Sleep spindle duration predicts decoding accuracy of activity-silent WM states

Having established that spindle duration during the nap predicted post-sleep WM performance beyond pre-nap performance, we next asked whether spindle physiology was also related to the neural readout of activity-silent WM representations after sleep. To address this, we tested whether individual differences in spindle duration predicted the fidelity of impulse-evoked decoding during the post-nap WM task.

To quantify neural readout of latent WM content, we decoded the remembered item orientation at impulse presentation using multivariate classification (10-class linear discriminant analysis; 8-fold cross-validation), predicting which of the 10 possible grating orientations each trial belonged to (see Methods). Both impulses were designed to probe the same types of latent WM representations; accordingly, decoding was performed separately for each impulse and then averaged to obtain a single, more stable estimate of impulse-evoked WM representation fidelity.

As anticipated, impulse presentation elicited a modest but robust increase in the decodability of orientation-specific information, with decoding accuracy reliably exceeding chance from 130–490 ms post-impulse (cluster-corrected p < 0.001; Figure 3B). Decoding accuracy was then averaged across this time window to yield a single value per participant. To obtain a task-relevant summary measure of spindle duration, we averaged mean spindle duration across the electrode cluster that showed a significant association with post-nap behavioural WM performance in the partial correlation analysis (Figure 3A). This empirically defined cluster was taken to reflect the spindle dynamics most relevant for subsequent WM processing.

Critically, participants with longer mean spindle duration over occipito-parietal sites exhibited higher impulse-evoked decoding accuracy (Spearman’s ρ = 0.46, p = 0.01; Figure 3C), indicating that inter-individual differences in spindle duration during the nap predicted the fidelity with which latent WM representations could be read out after sleep. Control analyses indicated that this relationship did not appear to be driven by overall sleep architecture. Neither total sleep proportion (Spearman’s ρ = 0.08, p = 0.7), nor the proportion of N2+N3 sleep (Spearman’s ρ = - 0.32, p = 0.09) was significantly associated with decoding accuracy. Similarly, these sleep-architecture measures were not significantly related to post-nap behavioural accuracy (total sleep proportion: Spearman’s ρ = 0.09, p = 0.6, N2 + N3 proportion: Spearman’s ρ = -0.20, p = 0.28).

Together, these findings demonstrate that specifically longer spindle duration during a daytime nap predicts post-sleep WM performance and neural readout of activity-silent WM content.

## Discussion

Building on the idea that activity-silent WM may be supported by transient changes in synaptic efficacy, we asked whether sleep spindles shape how readily latent WM representations can be accessed after sleep. Across 30 participants, longer mean spindle duration during NREM sleep over occipito-parietal sites was associated with better post-nap behavioural WM performance and, critically, with higher impulse-evoked decoding accuracy of item-specific (“silent”) WM representations. These relationships were selective: slow oscillation (SO) duration showed no corresponding associations, supporting the idea that spindle dynamics, and spindle duration in particular, are linked to the post-sleep fidelity with which latent WM content can be read out.

### Sleep spindles and post-sleep readout of activity-silent WM

Hybrid models of WM maintenance propose that recently encoded representations can be stored in transient synaptic states and reactivated through brief bursts of neural activity when needed (Mongillo et al., 2008; Panichello et al., 2024; Stokes, 2015). The impulse-perturbation (“pinging”) approach builds on this framework by perturbing the network during WM maintenance, eliciting impulse responses that carry information about latent memory content detectable with EEG decoding (Wolff et al., 2015, 2017). In the present study, impulse presentation during the post-nap WM task reliably reinstated orientation-specific information, and individual differences in decoding fidelity were predicted by spindle duration during the preceding nap. This extends the impulse-perturbation approach by linking variability in impulse-evoked WM readout to a well-characterised mediator of synaptic plasticity, i.e., sleep spindles (Dickey et al., 2021; Fernandez & Lüthi, 2020), and reveals a previously underexplored connection between spindle dynamics and latent WM states. Note that this association does not imply a one-to-one mapping between spindle duration and synaptic plasticity, or between impulse-evoked decoding and purely activity-silent storage. Nevertheless, it provides converging evidence that spindles, which have been closely linked to synaptic plasticity (Seibt et al., 2017), predict a WM readout designed to be sensitive to latent representations. This convergence makes spindle-mediated recalibration a plausible mechanism for post-sleep changes in WM accessibility.

A key finding of our study is the prominence of spindle duration as the spindle metric most closely associated with both post-sleep WM behaviour and impulse-evoked decoding accuracy. Rather than indexing sleep quantity per se, spindle duration may reflect the temporal window over which spindle-associated plasticity processes can act (Dittman & Regehr, 1998; Rosanova & Ulrich, 2005; Seibt et al., 2017). Longer spindles, containing more oscillatory cycles, may therefore provide extended opportunities for Ca²⁺-dependent synaptic recalibration. Such recalibration could manifest in several, not mutually exclusive, ways: selective strengthening of recently active connections, renormalisation of local circuits to improve dynamic range and signal-to-noise, or stabilisation of neuronal ensembles supporting latent WM representations. Any of these processes would be expected to yield a more robust and discriminable impulse-evoked “echo” of stored content after waking. Our observation that spindle duration predicted both behavioural performance and neural decoding fidelity is consistent with this interpretation, while leaving open which combination of synaptic recalibration and/or general network excitability properties give rise to the observed effect.

### Occipito-parietal topographies and trait-state considerations

The spindle-duration effects observed here align with accumulating evidence linking spindle characteristics to cognitive performance. While much classical work has focused on long-term memory consolidation and the coordination of spindles with slow oscillations and hippocampal ripples (Diekelmann & Born, 2010; Staresina, 2024), spindles have also been implicated in attentional and executive functions, and spindle duration has been associated with WM impairments in clinical risk populations (Mayeli et al., 2022). Our findings extend this work by tying spindle duration specifically to the post-sleep neural accessibility of latent WM representations – an aspect of WM function that is not captured by behaviour alone and that is theoretically grounded in short-term synaptic plasticity models.

Notwithstanding the regional ambiguity of scalp EEG, the occipito-parietal distribution of the spindle/WM association (Figure 3A) further accords with the visuo-spatial nature of the task and with emerging evidence that spindles over occipital and temporo-occipital cortex play a role in sleep-dependent plasticity within visual systems. For example, occipital spindle activity predicts overnight improvements in visual sequence learning (Lutz et al., 2021), and intensive visual training can selectively reorganise fast spindle timing over posterior sites (Gerván et al., 2025). Furthermore, a recent study – although not specifically looking at sleep spindles – has found that a daytime nap can recalibrate synaptic plasticity (Fehér et al., 2026). Together with these findings, the present results support the idea that spindle-related plasticity may operate locally within task-relevant cortical networks.

In our data, spindle duration correlated with post-nap performance but not with pre-nap accuracy. In fact, analytically controlling for pre-nap performance suggested that the post-sleep relationship could not be fully explained by stable individual differences. This pattern is compatible with a sleep-state contribution to WM function, but nevertheless warrants consideration of trait-like stability of spindle characteristics. Spindle measures have been linked to general cognitive ability (Fang et al., 2017), with meta-analytic work suggesting particularly robust trait associations for spindle amplitude and, to a lesser extent, density (Ujma, 2018). At the same time, night-to-night increases in spindle activity track learning-related memory improvements beyond baseline ability (Schabus et al., 2008), indicating that spindles can also reflect state-dependent plasticity. Taken together, these observations suggest that different spindle metrics may index partially separable trait and state components (Lustenberger et al., 2015). While our partial-correlation analyses are consistent with a sleep-dependent contribution, future within-subject and manipulation-based designs will be required to more clearly dissociate stable individual differences from sleep-induced changes in WM-relevant neural dynamics.

### Limitations and future directions

Impulse-evoked decoding provides only an indirect assay of latent WM states and cannot measure the synaptic variables posited by activity-silent models, such as presynaptic calcium dynamics, release probability or vesicle availability (Mongillo et al., 2008). Moreover, the present evidence is correlational. Spindle dynamics were not experimentally manipulated, and causal tests will require targeted interventions, for example closed-loop auditory stimulation designed to modulate spindle duration. Finally, recordings were obtained during a daytime nap; overnight studies will be an interesting future addition to more comprehensively explore interactions between spindles and slow oscillations.

Another important direction for future work concerns the role of pre-sleep task engagement. Participants performed a visual WM task prior to the nap, which was essential for indexing baseline, trait-like differences in WM ability, but may itself have influenced subsequent sleep physiology. Prior work has shown that engagement of specific cortical circuits within those same networks, particularly in terms of spindle density and amplitude (e.g., Clemens et al., 2005, 2006; Petzka et al., 2022; Tamaki et al., 2013; Thom & Staresina, 2025). Future studies incorporating baseline sleep sessions without prior task engagement will be important for determining whether the posterior spindle effects observed here reflect, at least in part, use-dependent modulation of spindle activity induced by the pre-nap WM task itself. Relatedly, the pre-sleep task may have created a particular demand for synaptic recalibration during sleep. Intensive engagement of visual WM circuits could lead to transient synaptic saturation, increasing the need for subsequent renormalisation or fine-tuning processes during NREM sleep. From this perspective, spindle-dependent recalibration might not only prepare cortical circuits for future WM performance in a generic sense, but may specifically counteract use-dependent changes incurred during prior task engagement. This idea resonates with a work on long-term memory showing that sleep restores the capacity for subsequent learning (March et al., 2023). Although these studies focus on episodic learning rather than WM, they are consistent with the possibility that spindle-related recalibration helps restore network-level dynamic range for subsequent information processing. Future studies should therefore test whether omitting a pre-sleep visual WM task, or replacing it with a task engaging different cortical systems, alters the relationship between spindle dynamics and post-sleep WM behaviour and neural decoding. Such designs, ideally combined with baseline sleep recordings, would help disentangle trait-like spindle characteristics from experience-dependent, circuit-specific modulation and clarify whether spindle-related effects on post-sleep WM depend on prior synaptic load within the relevant networks.

## Conclusion

The present findings provide evidence that the duration of sleep spindles may shape how effectively latent, presumably synaptic, WM traces can be reactivated after waking. In doing so, they extend sleep-dependent plasticity research beyond long-term systems consolidation and suggest that a restorative nap may not merely safeguard previous experiences, but can also prime cortical circuits for the encoding and high-fidelity readout of new information upon awakening.

## Acknowledgments

This project has received funding from the European Research Council (ERC) under the European Union’s Horizon 2020 (grant agreement no. 101001121) awarded to B.P.S..

## Author Contributions

Conceptualisation, S.A.W., E.G.A., and B.P.S.; methodology, S.A.W., E.G.A., and B.P.S.; formal analysis, S.A.W.; investigation, S.A.W.; writing – original draft, S.A.W.; writing – review & editing: E.G.A., B.P.S.; supervision: B.P.S.; funding acquisition: B.P.S.

## Data and Code Accessibility

Pre-processed data, and original code to support the conclusions of this study will be made publicly available upon publication on the Open Science Framework.

## Methods

### Participants

We recruited 34 healthy volunteers. Data from four participants were excluded because of technical problems during recording (n = 2), poor task accuracy (< 70%; n = 1), or early termination due to illness (n = 1), leaving a final sample of 30 participants (mean age 24.8 ± 5.9 years; 18 female; 28 right-handed), in keeping with previous WM impulse-perturbation work (Kandemir, Wilhelm, et al., 2024; Kandemir, Wolff, et al., 2024; Wolff et al., 2015, 2017; Wolff, Jochim, et al., 2020; Wolff, Kandemir, et al., 2020). Prior to laboratory testing, volunteers completed an online screening questionnaire. Individuals were excluded if they had worked night shifts in the preceding year, reported neurological, psychiatric, or sleep disorders, or used sleep-altering medication. All invited participants scored < 5 on the Pittsburgh Sleep Quality Index (PSQI; Buysse et al., 1989). Participants additionally completed a ∼15-minute online version of the working-memory task on Pavlovia, and only those achieving ≥ 65% accuracy on the experimental blocks were invited to the laboratory. The task was identical to the post-nap working memory task (Task 2), and served to verify task comprehension (particularly the clockwise/counterclockwise judgments) before invitation to the laboratory.

Participants were instructed to avoid alcohol on the day prior to the study, to wake approximately one hour earlier than usual on the study day, and to abstain from caOeine. All participants provided written informed consent and received £70 for participation. The study was approved by the Oxford University Central Research Ethics Committee (CUREC #R90055/RE001).

### Procedure

Each laboratory session lasted from 10:00 to 17:00. Following consent, participants washed and towel-dried their hair, after which a 64-channel EEG cap was applied, including two electrooculography (EOG), two electromyography (EMG), and bilateral mastoid electrodes, leaving 58 scalp EEG channels.

At approximately 11:00, participants completed the pre-nap working-memory task (Task 1; 45–60 min). They then lay in a darkened room for a 90-min nap opportunity. After awakening, participants were given ∼15 min to eat and fully wake before beginning the post-nap working-memory task (Task 2), which comprised two 1.5-h blocks separated by a 15-min break during which the EEG amplifier was disconnected to allow movement.

Sessions ended with cap removal and hair washing, and participants completed the study at approximately 17:00.

### Task

Tasks were programmed in MATLAB (MathWorks) using Psychtoolbox (Brainard, 1997) and presented on a Dell S2522HG 24.5-inch Fast IPS LED-backlit LCD monitor (1920 × 1080 px, refresh rate = 60Hz) driven by an 2500 Intel UHD Graphics 770 GPU. Participants viewed the screen from a distance of ∼60 cm. The background was grey (RGB = 128,128,128), and a central black fixation dot was present except during cue presentation.

Memory items and probes were circular gratings (diameter = 6.69°, spatial frequency = 0.65 cycles/°, 20% contrast). On each trial, orientations were drawn without replacement from ten predefined angles (5.25°, 23.75°, 42.25°, 60.75°, 79.25°, 97.75°, 116.25°, 134.75°, 153.25°, 171.75°). Probe orientations differed from the target by ±10°, ±16°, ±24°, ±26°, ±32°, or ±40°, following Wolff et al. (2017). Impulse (“ping”) stimuli consisted of a centrally presented white disk (diameter = 13.38°).

All stimuli were presented centrally, and responses were collected using a standard keyboard.

#### Task 1 (Pre-nap)

Fixation intervals were jittered by ±100 ms. Each trial began with 900 ms fixation, followed by two memory items presented sequentially for 200 ms each and separated by 900 ms fixation. After the second item, fixation remained for 900 ms before a retro-cue indicated which item would be tested. Nine hundred milliseconds later, the probe appeared for 250 ms. Participants had up to 1500 ms to respond, pressing “c” for clockwise and “m” for counter-clockwise rotations relative to the cued item.

Feedback (“:)” or “:(”) was presented immediately after the response. If no response was made within 1500 ms, the trial advanced automatically. The session comprised 380 trials across 19 blocks of 20 trials, with self-paced breaks between blocks.

#### Task 2 (Post-nap)

Fixation jittering and encoding were identical to Task 1. After encoding, a cue (“1” or “2”) indicated whether the first or second encoded item should be reported first. Following 900 ms fixation, a 100-ms impulse was presented. Five hundred milliseconds later, the first probe appeared for 250 ms, and participants had up to 1500 ms to respond. If they responded early, fixation filled the remainder of the interval to equate timing across trials.

Feedback was presented for 200 ms, followed by 900 ms fixation, a second 100-ms impulse, and after 500 ms, a second probe. The second response followed identical rules. The task comprised 960 trials arranged into 40 blocks of 24 trials, divided into two sessions separated by a 15-min break.

### EEG data acquisition

EEG was recorded with a 64-channel Brain Products system (Brain Products UK) at 500 Hz. Electrode placement followed the international 10–20 system, with FCz as ground and AFz as online reference. Impedances were maintained below 10 kΩ. EMG electrodes were placed under the chin, and mastoid and EMG electrodes were secured with medical tape. The cap was wrapped with bandage gauze to maintain stability during sleep.

### Sleep scoring

Sleep was scored in 30-s epochs using two automated classifiers (SomnoBot; YASA (Vallat & Walker, 2021). EEG was re-referenced to linked mastoids, and hypnograms from both tools were imported into MATLAB. A trained researcher reviewed all epochs in which classifiers disagreed and resolved discrepancies according to American Academy of Sleep Medicine criteria (Berry et al., 2017).

### Sleep EEG preprocessing and event detection

Sleep EEG was processed using FieldTrip (Oostenveld et al., 2011). Bad channels were interpolated using spherical splines, retaining 58 scalp electrodes which were then re-referenced to the common average. Artefactual segments were identified using MATLAB’s isoutlier function based on extreme amplitude, mean absolute deviation, or high-frequency power.

Spindles and slow oscillations (SOs) were detected during artefact-free N2 and N3 sleep using custom code adapted from Ngo et al. (2020). Spindles were identified after band-pass filtering at 12–16 Hz and smoothing with a 200-ms moving average; events exceeding mean + 1.25 SD in amplitude and lasting 0.5–3 s were classified as spindles. Spindle amplitude was defined as the maximum of the envelope, spindle density as events per minute, and spindle duration as onset-to-offset length.

SOs were detected after filtering at 0.3–1.25 Hz. Zero crossings defined candidate events, which were retained if lasting 0.8–2 s and exceeding amplitude thresholds (1.25 SD for trough and trough-to-peak). SO amplitude was defined as trough voltage, density as events per minute, and duration as full event length.

### Task EEG preprocessing

Task EEG was processed in FieldTrip. Independent component analysis was applied using ft_componentanalysis, and components reflecting ocular artefacts were removed following visual inspection. Bad channels were interpolated, data re-referenced to the common average, filtered between 0.1–40 Hz, and epoched from −0.5 to 0.9 s relative to each event. Residual artefacts were removed using ft_rejectvisual, discarding epochs with z-scores > 2.

### Decoding Analysis

All decoding was performed in MATLAB using MVPA-Light (Treder, 2020). A 10-class linear discriminant analysis classifier was trained to predict the orientation of the memorised items.

### Decoding Time Course

Data were baseline-corrected (−0.2 to 0 s), down-sampled to 100 Hz, restricted to −0.1 to 0.5 s, and smoothed with a 20-ms moving average. “Super-trials” were created by averaging five trials, and class sizes were equalised by undersampling. Eight-fold cross-validation was repeated 50 times per time point, and accuracies were averaged across repetitions.

### Decoding Topography

Searchlight decoding was performed using each electrode and its neighbours as features. Decoding was repeated across time, and topographies were obtained by averaging accuracies across significant time points from the main time-course analysis.

### Statistics

The chance level was predefined as 10% for the 10-class LDA classifier. True decoding accuracy was tested against this chance level at each time point using a one-sided dependent-samples t-test across participants. We additionally verified the results using subject-specific null estimates generated by repeating the decoding analysis 250 times with shuffled labels. As this yielded the same pattern of results, we report the analysis using the predefined 10% chance level.

Multiple comparisons across time were controlled using cluster-based permutation testing in FieldTrip with a cluster-forming threshold of *p* < 0.05 and cluster-level significance assessed using 10,000 Monte Carlo randomisations.

Associations between decoding accuracy and sleep oscillation measures were assessed using Spearman correlations, with cluster-based correction across electrodes where appropriate. Statistical significance was set at p < 0.05.

For electrode-wide topographical analyses, correlations were computed at each electrode and corrected for multiple comparisons across electrodes using cluster-based permutation testing in FieldTrip based on neighbouring electrodes. Spatial clusters were formed at *p* < 0.05, and cluster-level significance was assessed using 10,000 Monte Carlo randomisations. This procedure was used for both the supplementary topographical analyses and partial-correlation analysis in the main text. All tests were two-sided unless stated otherwise.

