## Supplementary Materials for "Sleep spindles enhance latent working memory representations"

**SUPPLEMENTARY INFORMATION**

**Supplementary Table 1:** Summary of sleep stages

| **Metric** | **Mean (SD)** | **Min** | **Max** |
| --- | --- | --- | --- |
| Time N1 (mins) | 11.5 (6.2) | 3 | 30 |
| Time N2 (mins) | 36.7 (10.5) | 18.5 | 60 |
| Time N3 (mins) | 11.1 (11.9) | 0 | 43 |
| Time N2/N3 (mins) | 47.9 (11.8) | 24 | 67.5 |
| Time REM (mins) | 6.8 (8.3) | 0 | 24.5 |
| Total Sleep Time (mins) | 66.2 (14.8) | 33.5 | 85.5 |
| Proportion N1 | 0.17 (0.09) | 0.05 | 0.43 |
| Proportion N2 | 0.57 (1.5) | 0.3 | 0.88 |
| Proportion N3 | 0.16 (0.16) | 0 | 0.58 |
| Proportion N2/N3 | 0.73 (0.13) | 0.41 | 0.92 |
| Proportion REM | 0.09 (0.1) | 0 | 0.35 |

**Supplementary Table 2:** Summary of sleep oscillation characteristics

| **Event** | **Electrode** | **Metric** | **Mean (SD)** | **Min** | **Max** |
| --- | --- | --- | --- | --- | --- |
| **Spindle** | Cz | Density | 4.5 (0.5) | 3.4 | 5.5 |
|  |  | Amplitude | 6.6(2.5) | 0.5 | 13.6 |
|  |  | Duration | 0.83 (0.06) | 0.7 | 0.92 |
| **Slow oscillation** | Fz | Density | 2.2 (0.6) | 0.9 | 3.2 |
|  |  | Amplitude | 29.8(9.7) | 13.6 | 52.5 |
|  |  | Duration | 1.5 (0.04) | 1.4 | 1.6 |

**Supplementary Figure 3:**

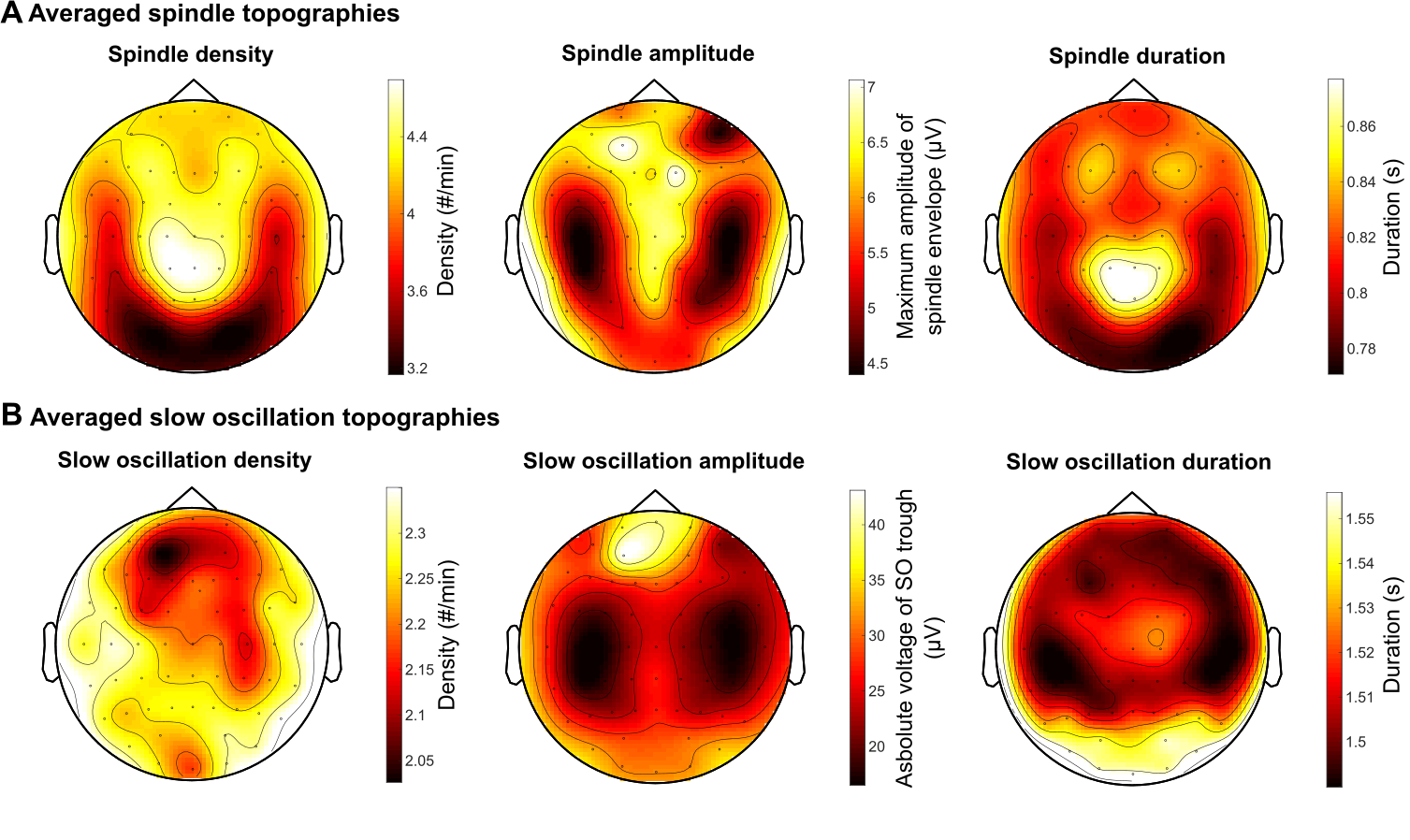

Figure: **Topographical overview of spindle and slow oscillation characteristics averaged across participants. A** Average sleep spindle topographies. Left to right: density, power and duration. **B** Average slow oscillation topographies. Left to right: density, amplitude and duration. Colour bars indicate the metric scale of each sub-plot.

**Supplementary Figure 4:**

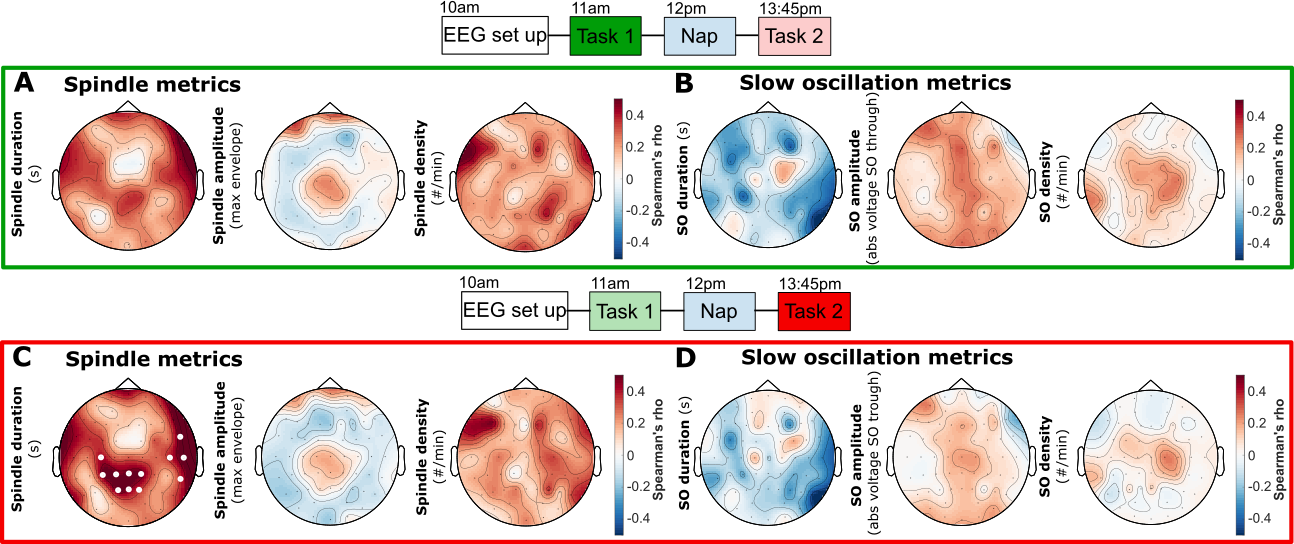

**Control analyses for spindle and slow oscillation metrics across tasks.** Scalp topographies show Spearman correlations between behavioural WM accuracy and sleep-oscillation metrics computed at each electrode, with cluster-based correction for multiple comparisons across electrodes. **A** Scalp topography of correlation between pre-nap Task 1 accuracy and spindle duration (*left*)*,* spindle amplitude (*middle*) and spindle density (*right*). No significant clusters for any metric (all *p* > 0.05). **B** Corresponding correlations between pre-nap Task 1 accuracy and SO metrics: duration (*left*), amplitude (*middle*), and density (*right*). No significant clusters for any metrics (all *p* > 0.05). **C** Scalp topographies of correlation between post-nap Task 2 accuracy and the same spindle metrics. Significant correlation between spindle duration and post-nap WM accuracy across two clusters (*central:* cluster-corrected *p* = 0.025; *right temporal*: cluster-corrected *p* = 0.045). No significant clusters for spindle amplitude and spindle density (all *p* > 0.05). **D** Corresponding correlations between post-nap Task 2 accuracy and SO metrics. No significant clusters emerged (all *p* > 0.05).
